# Inference of Gorilla demographic and selective history from whole genome sequence data

**DOI:** 10.1101/009191

**Authors:** Kimberly F. McManus, Joanna L. Kelley, Shiya Song, Krishna Veeramah, August E. Woerner, Laurie S. Stevison, Oliver A. Ryder, Jeffrey M. Kidd, Jeffrey D. Wall, Carlos D. Bustamante, Michael F. Hammer

## Abstract

While population-level genomic sequence data have been gathered extensively for humans, similar data from our closest living relatives are just beginning to emerge. Examination of genomic variation within great apes offers many opportunities to increase our understanding of the forces that have differentially shaped the evolutionary history of hominid taxa. Here, we expand upon the work of the Great Ape Genome Project by analyzing medium to high coverage whole genome sequences from 14 western lowland gorillas (*Gorilla gorilla gorilla*), 2 eastern lowland gorillas (*G. beringei graueri*), and a single Cross River individual (*G. gorilla diehli*). We infer that the ancestors of western and eastern lowland gorillas diverged from a common ancestor ∼261 thousand years ago (kya), and that the ancestors of the Cross River population diverged from the western lowland gorilla lineage ∼68 kya. Using a diffusion approximation approach to model the genome-wide site frequency spectrum, we infer a history of western lowland gorillas that includes an ancestral population expansion of ∼1.4-fold around ∼970 kya and a recent ∼5.6-fold contraction in population size ∼23 kya. The latter may correspond to a major reduction in African equatorial forests around the Last Glacial Maximum. We also analyze patterns of variation among western lowland gorillas to identify several genomic regions with strong signatures of recent selective sweeps. We find that processes related to taste, pancreatic and saliva secretion, sodium ion transmembrane transport, and cardiac muscle function are overrepresented in genomic regions predicted to have experienced recent positive selection.

## Introduction

The *Gorilla* genus consists of two morphologically distinguishable species, western (*Gorilla gorilla*) and eastern (*Gorilla beringei*) gorillas (Grubb et al. 2003), each of which is divided into two recognized subspecies. Eastern gorilla populations occur in lowlands and highlands in the Democratic Republic of Congo, Uganda and Rwanda; while western gorilla populations reside primarily in Cameroon, Equatorial Guinea, Gabon, Congo, and the Central African Republic (Thalmann et al. 2007). Western gorillas include western lowland gorillas (*Gorilla gorilla gorilla*), the subspecies with the largest population size (and the main focus of this study), and Cross River gorillas (*G. gorilla diehli*), of which only a few hundred individuals remain. Eastern gorillas are composed of eastern lowland gorillas (*G. beringei graueri*) and mountain gorillas (*G. beringei beringei*), which are found today in only two small isolated subpopulations.

Gorillas are the largest extant nonhuman primate, with male and female western lowland gorilla body weights averaging 170 and 71 kg, respectively (Smith and Jungers 1997). Gorillas also demonstrate the largest sexual dimorphism in body size of any of the apes. This is likely related to their mating system (Plavcan 2001), where gorillas exhibit a polygynous structure in which a single dominant male largely controls access to reproduction with a number of adult females. Among the apes, gorillas also demonstrate an unusual diet and digestive anatomy. For example, field studies indicate that the western gorilla diet comprises as many as 230 plant parts from 180 plant species (Rothman et al. 2006). While consuming extremely diverse and large quantities of terrestrial vegetation throughout the year, western gorillas will also regularly eat fruit when it is available (Rogers et al. 2004; Doran-Sheehy et al. 2009). Given their large body size and strictly herbivorous/frugivorous diet, it is not surprising that the gorilla gut anatomy has evolved a distinctive digestive anatomy and physiology. Mainly as a result of the large capacity for microbial fermentation in a large pouched colon, gorillas gut anatomy allows for energy gain through the absorption of volatile fatty acids and microbial protein (Stevens and Hume 1995).

Research on demographic events and selective pressures experienced by gorillas may provide insights to the evolutionary forces that have uniquely influenced patterns of gorilla morphological and genetic variation. Both species of gorilla are considered threatened on the IUCN Red List of Threatened Species (IUCN 2013); western gorillas are classified as critically endangered and eastern gorillas are classified as endangered. Recent census estimates indicate a rapid recent population size contraction in gorillas due to multiple factors including: outbreaks of the Ebola virus, the bushmeat trade, habitat loss and fragmentation (Anthony et al. 2007; Le Gouar et al. 2009; Walsh et al. 2003).

In this study, we use a coalescent approach to infer divergence times, migration rates, and effective population sizes based on medium to high coverage whole genome data from three gorilla subspecies: western lowland, Cross River and eastern lowland. We use a diffusion approximation approach to infer temporal changes in western lowland gorilla effective population size, conduct a genome scan for positive selection to identify signatures of recently completed selective sweeps, and investigate the nature of genetic changes within those regions to identify putative targets of selection. Additionally, we analyze the overall distribution of fitness effects for polymorphic sites in western lowland gorillas and the proportion of substitutions compared to humans that have been driven by adaptive evolution versus random genetic drift.

## Results

### Gorilla population structure

Whole genome sequence data from 17 gorillas, including 14 western lowland gorillas, 2 eastern lowland gorillas, and 1 Cross River gorilla were aligned to the gorGor3 (Ensembl release 62) reference genome and processed with filtering as previously described (Prado-Martinez et al. 2013, Table S1). We limited our analysis to samples without evidence of inter and intra-species sequence contamination and characterized patterns of genetic variation based on SNP genotypes obtained using the GATK Unified Genotyper, limited to sites with at least eight-fold (8x) coverage in all samples. Across the autosomes, we observe that eastern lowland gorillas have the lowest heterozygosity (5.62—5.69 × 10^−4^) of all the groups studied here followed by the single Cross River sample (9.09 × 10^−4^) and 14 western lowland gorillas (1.2—1.6 × 10^−3^) (**Figure S1**). We used principal components analysis (PCA) (Patterson et al. 2006) and ADMIXTURE (Alexander et al. 2009) to further explore relationships among the samples. As previously observed (Prado-Martinez et al. 2013), when considering all samples together, PC1 shows clear separation of eastern and western gorillas with western lowland and Cross River gorillas arrayed along PC2 (**Figure S2**). PCA performed on only the western lowland gorilla samples does not reveal clear population clusters, although the individuals are somewhat ordered by sample geography (**Figure S3**). Results from ADMIXTURE, a model-based clustering algorithm allowing for mixed ancestry, support the existence of two clusters dividing eastern and western lowland gorillas (**Figure S4**). When applied to only the 14 western lowland gorilla samples, we observe that K=1 has the lowest cross-validation (CV) error (**Figure S5**). Moreover, the appearance of a cline at K=2 suggests a poor fit of the data to an admixture model with discrete sources and PCA does support substructure in the form of a cline in western lowland gorillas (**Figure S3**).

### Relationship between Western Lowland and Eastern Lowland Gorillas

We applied Generalized Phylogenetic Coalescent Sampler (G-PhoCS), a Bayesian coalescent-based approach, to infer ancestral population sizes, divergence times, and migration rates (Gronau et al. 2011) amongst the three gorilla subspecies. This inference is based on genealogies inferred at many independent and neutrally evolving loci across the autosomal genome. To avoid bias caused by the alleles represented in the reference genome, which is derived from a western gorilla, we used BSNP (Gronau et al. 2011), a reference-genome–free Bayesian genotype inference algorithm, to perform variant calling separately for each sample. Based on the BSNP output, we produced diploid sequence alignments of two eastern lowland gorillas, nine western lowland gorillas, one Cross River gorilla, and the human reference genome at 25,573 “neutral loci” with size approximately 1 kilobase (kb) and an interlocus distance of approximately 50 kb. The neutral loci were chosen based on the positions of putatively neutral loci previously utilized for humans (Gronau et al. 2011); loci intersecting with recent transposable elements, exons of protein-coding genes, and segmental duplications in the gorilla genome were removed.

For many of the analyses presented here, we used a four-population phylogeny as inferred by TreeMix (Pickrell and Pritchard 2012) and in agreement with our ADMIXTURE results (**Figures S6** and **S7**), with eastern and western gorilla ancestors separating first, followed by western lowland and Cross River gorilla (Figure 1). We first evaluated four alternative scenarios, having either no migration between any gorillas (Figure 1, scenario 1) or bi-directional migration between any two gorilla species (Figure 1, scenarios 2, 3, and 4). In G-PhoCS, migration is modeled as migration bands of constant migration rate between two lineages over the entire time period of their existence. We utilized several combinations of western lowland gorilla samples, always including two eastern lowland gorillas, two western lowland gorillas, one Cross River gorilla, and one human. We initially ran G-PhoCS for 50,000 iterations and monitored convergence using Tracer (Rambaut et al. 2013). Estimates of population split times are sensitive to model assumptions, particularly migration. Our G-PhoCS analysis finds no evidence of migration events between western lowland and Cross River gorillas (**Figure S8**, scenario 2). We do observe evidence of migration from western lowland gorilla to eastern lowland gorilla with mean total migration rate 0.3 (95% CI: 0.240—0.356), equivalent to 0.37 migrants per generation (95% CI: 0.312—0.433) (**Figure S8**, scenario 4). We also observe a small signal of migration from Cross River gorilla to eastern lowland gorilla (**Figure S8**, scenario 3); however, 50,000 iterations were not sufficient for convergence. To further explore these results, we tested a scenario with two migration bands: one from western lowland to eastern lowland gorilla and another from Cross River to eastern lowland gorilla (Figure 1, scenario 5), and extended the number of iterations to 300,000 to allow the posterior estimates to fully converge (**Figure S9**). Setting an additional migration band from Cross River to eastern lowland gorilla makes little difference because migration from western lowland to eastern lowland gorilla has the strongest migration signal (**Figures S10-S13**). The estimated migration rate from Cross River to eastern lowland gorilla is 0.004 (95% CI: 0.000—0.018), equivalent to 0.019 migrants per generation (95% CI: 0.000—0.071). By using this setting (Figure 1, scenario 5), we estimate the split time between western lowland gorilla and Cross River gorilla to be ∼68 thousand years ago, and the split time between eastern lowland and western ancestral gorilla to be ∼261 thousand years ago when assuming a human and gorilla divergence time of 12 million years ago (Scally et al. 2012) (Table 1). We also observed a decrease of western gorilla population size and a decrease of eastern gorilla population size after their initial split and a six-fold difference between current eastern and western gorilla population size. The relative population sizes of the gorilla populations are rather robust to the chronological human/gorilla split time used for calibration, though the actual estimated size and chronological date of the split times are sensitive to the split time assumptions as many calculations are pegged to the calibration date (Table 1).

**Figure 1.**
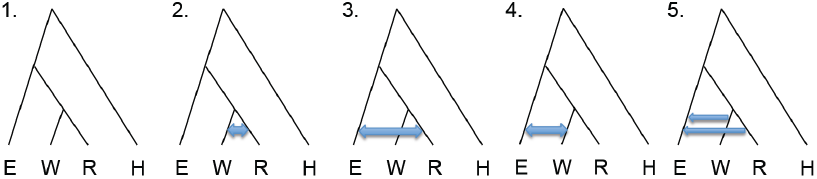
Phylogeny and explored migration bands for G-PhoCS analysis. We used the indicated phylogeny for eastern lowland (E), western lowland (W), Cross River (R) gorilla species and human (H), and tested the indicated migration scenarios. Scenario 1: no migration. Scenario 2: bi-directional migration between western and Cross River gorilla. Scenario 3: bi-directional migration between Cross River and eastern gorilla. Scenario 4: bi-directional migration between western and eastern gorilla. Scenario 5: migration from western to eastern gorilla and from Cross River to eastern gorilla.

**Table 1.** Gorilla population history estimates. Population history estimates by using G-PhoCS when assuming a range of human-gorilla divergence time (8, 10, and 12 Mya). We assumed migration events from western lowland to eastern lowland gorilla and from Cross River to eastern lowland gorilla (Figure 1 scenario 5). Values in parentheses correspond to 95% credible intervals.

|  | Human-Gorilla Divergence Time (Mya) |  |  |
| --- | --- | --- | --- |
|  | 8 | 10 | 12 |
| **Mutation rate per generation without CpG ($\times 10^{-8}$)** | 1.461<br>(1.456-1.466) | 1.169<br>(1.165-1.173) | 0.974<br>(0.970-0.978) |
| **Eastern Gorilla population size ($\times 10^3$)** | 2.853<br>(2.755-2.956) | 3.566<br>(3.443 -3.696) | 4.280<br>(4.132-4.435) |
| **Western Gorilla population size ($\times 10^3$)** | 16.774<br>(13.114 -21.439) | 20.967<br>(16.393-26.798) | 25.161<br>(19.672-32.158) |
| **Cross River Gorilla population size ($\times 10^3$)** | 2.054<br>(2.352-2.755) | 2.567<br>(2.940-3.443) | 3.080<br>(3.529-4.132) |
| **Western – Cross River ancestral population size ($\times 10^3$)** | 20.462<br>(17.294-24.191) | 25.578<br>(21.617-30239) | 30.693<br>(25.940-36.287) |
| **Gorilla ancestral population size ($\times 10^3$)** | 26.500<br>(25.829-26.965) | 33.126<br>(32.286-33.706) | 39.751<br>(38.743-40.447) |
| **Human-Gorilla ancestral population size ($\times 10^3$)** | 45.472<br>(44.349-46.608) | 56.840<br>(55.437-58.259) | 68.208<br>(66.524-69.911) |
| **Western – Cross River split time (Mya)** | 0.046<br>(0.038-0.056) | 0.057<br>(0.048-0.070) | 0.068<br>(0.057-0.084) |
| **Eastern – Western-Cross River ancestral split time (Mya)** | 0.174<br>(0.161-0.194) | 0.218<br>(0.201-0.243) | 0.261<br>(0.242-0.292) |

### Western Gorilla Demographic Inference

We additionally inferred the fine-scale population history of western lowland gorillas using the genome-wide site frequency spectrum (SFS) obtained from 14 individuals (Gutenkunst et al. 2009). We utilized a diffusion approximation for demographic inference (*∂α∂i*) on the unfolded SFS based on 4,554,752 SNPs only considering sites where all samples had at least 8x coverage.

Variants were polarized to ancestral and derived alleles based on human out-group sequences, and we implemented a context-dependent correction for ancestral misidentification (Hernandez et al. 2007). Five demographic models were fit using *∂α∂i* and inferring the best-fit demographic model requires us to assess whether the improvement in fit afforded by additional parameters needed in more complex models are justified (Table 2). Our results suggest an ancient expansion followed by a more recent drastic, nearly three-fold, population contraction is the best model for the data. Specifically, assuming a mutation rate of 1.1 × 10^−8^ per base pair per generation (Roach et al. 2010) and generation time of 19 years (Langergraber et al. 2012), the best-fit model is a three-epoch model that has an ancestral effective population size of 31,800 (95% CI: 30,690—32,582) (Table 2). While the bottleneck followed by exponential growth model and the three-epochs models have similar fits, the three-epochs has the best fit; moreover, the model selection is robust when SNPs are thinned to 100kb. The first size change event occurred 969,000 years ago (95% CI: 764,074—1,221,403) and increased the effective population size to 44,200 (95% CI: 42,424—46,403) individuals. The second size change event occurred 22,800 years ago (95% CI: 16,457—30,178) and decreased the effective population size to 7,900 (95% CI: 6,433—9,240) individuals (Figure 2).

**Figure 2.**
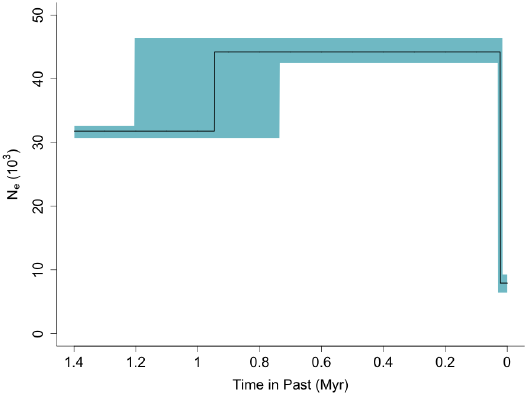
Inferred best-fit demographic model of western lowland gorillas. Shading represents confidence intervals determined by bootstrapping. Fitted parameters are depicted assuming a mutation rate of 1.1 × 10^−8^ per base pair per generation and a generation time of 19 years.

**Table 2.** Demographic model inference results from the five demographic models tested in #x1D74F;#x1D482;#x1D74F;#x1D48A; to fit the 8x unfolded SFS for western lowland gorillas. Gray line contains program parameter output, and white line contains conversion into years. With P1 first population size change, T1 length of bottleneck, P2 second size change, and T2 time of second size change. For the conversion, a mutation rate of 1.1e-8 mutations per base pair per generation and a 19-year generation time were used. The total number of callable sites is 812,645,853.

| Demographic Model | Theta / Ancestral Pop Size | P1 | T1 | P2 | T2 | Log-likelihood | AIC |
| --- | --- | --- | --- | --- | --- | --- | --- |
| Standard Neutral | 1,167,204 |  |  |  |  | -60,420 | 120,840 |
|  | 32,643 |  |  |  |  |  |  |
| Exponential Growth | 1,299,805 | 0.09 | 0.009 |  |  | -6,222 | 12,448 |
|  | 36,352 | 3,272 | 12,432 |  |  |  |  |
| Bottleneck, then Exponential Growth | 1,181,405 | 39.54 | 0.33 | 0.32 |  | -578 | 1,162 |
|  | 33,040 | 1,306,416 | 10,903 | 401,771 |  |  |  |
| Two Epochs | 1,297,300 | 3.4e-13 | 1.2e-14 |  |  | -5,654 | 11,312 |
|  | 36,282 | 0 | 0 |  |  |  |  |
| Three Epochs | 1,136,249 | 1.391 | 0.785 | 0.249 | 0.019 | -473 | 954 |
|  | 31,777 | 44,190 | 946,129 | 7,905 | 22,842 |  |  |

Nsubuga et al. (2010) and Fünfstück et al. (2014) found evidence for multiple population clusters within western lowland gorillas utilizing data from simple sequence repeats (SSR; microsatellite) variation. Though our ADMIXTURE results on our data find the lowest CV error with a one-population model, we also inferred demography separately for individuals on either side of the putative cline (**Table S2**). Both sample sets yield very similar demographic inferences compared to those obtained from the combined set of 14 individuals.

### Selection in Western Lowland Gorillas Identifying Selective Sweeps

We employed a composite likelihood approach (SweeD, Pavlidis et al. 2013) to scan for genomic regions showing signs of recent selective sweeps (Nielsen et al. 2005). The method compares the regional SFS to the background SFS to calculate a composite likelihood ratio (CLR), which indicates the likelihood of a sweep at a specific genomic region (in 100kb windows).

Significance was determined by comparisons to neutral regions (without selection) simulated in ms (Hudson 2002) with the inferred three-epoch demography. Genomic windows were compared to simulated regions with similar estimated recombination rate and percent of sequence masked. For the autosomes, these analyses identified 273 windows of size 100kb with p < 10^−3^. With a more stringent p-value cut-off a subset of the windows are identified: 111 windows of size 100kb with p < 10^−4^ (**Table S3**). A total of 50 windows had a p-value < 10^−5^, indicating that the CLR of these windows surpassed every CLR obtained from the simulated neutral distribution. As some of the 50 windows were adjacent to each other, these correspond to 43 distinct regions where the observed CLR value exceeded that obtained from 100,000 neutral simulations.

The region with the largest CLR in the western lowland gorilla genome is located on chromosome 5 (Figure 3). This region consists of four adjacent 100kb windows with p-value < 10^−5^ (chr5: 122,465,120 – 122,864,624). There are several genomic features in this top-scoring 400 kb region, including the protein coding genes *CTNNA1*, *SIL1*, and *MATR3* as well as other non-coding features, including *5S rRNA*, *U6* snRNA, and *SNORA74*. The region contains three non-synonymous SNPs which pass the quality filtering but do not have coverage to pass the 8x depth filter: one coding change in *CTNNA1*, a cadherin-associated protein, and two in *SIL1*, a nucleotide exchange factor which interacts with heat-shock protein-70. We also note that this region is directly upstream of *SLC23A1,* a vitamin C transporter and *PAIP2*, a repressor of polyadenylate-binding protein PABP1. *PAIP2* acts as part of innate defense against cytomegalovirus (CMV) (McKinney et al. 2013), which has been detected in wild gorilla populations (Leendertz et al. 2009).

**Figure 3.**
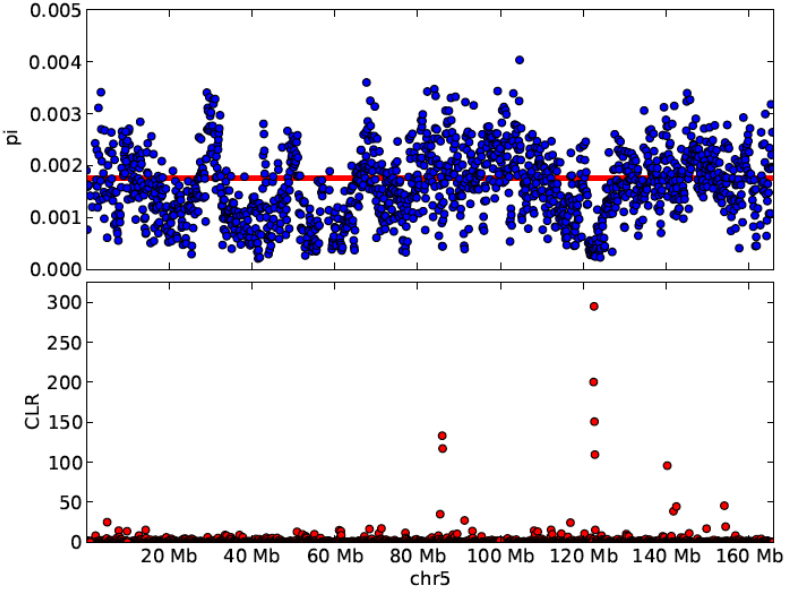
Signatures of selective sweeps in western lowland gorillas. Sequence diversity (pi) and the composite likelihood ratio (CLR) from the test for selective sweeps plotted for chromosome 5. The red line indicates the genome-wide average value for pi from 14 western lowland gorillas. Only windows with at least 10 kb of passing the 8x mask criteria are shown.

Another region identified (at p-value <10^−4^) contains several genes involved in taste reception. While in the 8x dataset, there is one nonsynonymous SNP in *TAS2R20*, our full SNP data set contains segregating non-synonymous changes in three of the taste receptors, including one change in TAS2R50 (derived allele frequency (DAF) = 93%), three in TAS2R20 (DAFs = 89%, 11% and 7%) and two in TAS2R19 (DAFs = 7% and 4%).

We conducted a gene ontology enrichment analysis of all regions with p < 10^−3^ using the Bioconductor package, topGO (Alexa and Rahnenfuhrer 2010) to identify gene pathways subjected to recent selective sweeps in western lowland gorillas. Using the elimination method with Fisher’s exact test, we identified 16 enriched GO categories (p < 0.01) (**Table S4**). The term with the lowest p-value is sodium ion transmembrane transport (GO:0035725, p=0.00039) and terms related to taste, pancreatic and saliva secretion, cardiac muscle cell function, and several others were identified.

### Distribution of Fitness Effects

We estimated *α*, the fraction of non-synonymous mutations to reach fixation due to adaptive evolution, through the method outlined in Keightley and Eyre-Walker (2012). This method utilizes the synonymous and nonsynonymous SFS, as well as divergence relative to an outgroup, to simultaneously infer demography and the distribution of fitness effects assuming a gamma distribution. Using the human reference genome as an outgroup we estimate *α* for western lowland gorillas to be 1.4% (95% CI: −11.6% – 11.0%). The distribution of fitness effects has a shape parameter of 0.152 and mean *N*_*e*_*s* of 3076. This distribution is leptokurtic, with a strong peak near zero and a long negative tail that extends to lethality, indicating that the vast majority of fixed and segregating non-synonymous variants are nearly neutral.

## Discussion

### Relationship between western lowland, Cross River and eastern lowland gorillas

Several groups have previously estimated split times and population sizes for western and eastern gorillas (Ackermann and Bishop 2010, Prado-Martinez et al. 2013, Scally et al. 2012, Thalmann et al. 2007, Thalmann et al. 2011) (**Table S5**). These studies make use of disparate data sets and modeling assumptions, particularly in terms of the treatment of gene flow subsequent to initial population separations. Based on 8 microsatellites, Thalmann et al. (2011) estimate that the separation of Cross River and western lowland gorilla populations occurred 17.8 kya, followed by a comparatively high level of gene flow. On the other hand, Prado-Martinez et al. (2013) estimated this population divergence time at 114 kya based on a modified PSMC approach (Note: the above mentioned values have been adjusted to match the mutation rate used in this manuscript where appropriate). The random phasing procedure applied in the modified PSMC approach may not be appropriate for such recent population split times (Prado-Martinez et al. 2013). Our estimate of 68 kya for the Cross River–western lowland split is intermediate between the two above estimates; however, we do not find support for gene flow between these two groups in our G-PhoCS analysis.

We estimate that the separation of eastern gorillas from the western lowland/Cross River ancestor occurred 261 kya, with subsequent migration from both western lowland and Cross River populations to the eastern gorillas. This value is similar to the 214 ky split time inferred by the modified PSMC approach. Scally et al. (2012), based on a model of symmetric migration, estimated a separation time of 429 kya. Mailund et al. (2012) arrives at a broadly similar estimate based on a coal-HMM, and estimates gene flow continuing until 150 kya. We note that our analysis indicates that the migration direction was from western lowland and Cross River to eastern gorillas, with more migrants coming from western lowland than from Cross River gorilla. However, Thalmann et al. (2007) find evidence for gene flow from eastern to western gorillas.

Alternatively, Ackermann and Bishop (2010) find support for a western to eastern gene flow in morphological and molecular data. The D-statistics (Green et al. 2010; Durand et al. 2011) calculated in Prado-Martinez et al. (2013) suggest that Cross River gorillas are genetically closer to eastern gorillas than western lowland gorillas are to eastern gorillas, which would not be predicted by the migration rates we infer. We further explored this apparent contradiction by calculating D-statistics for additional samples from Prado-Martinez et al. (2013) and using variants identified by BSNP based on mapping to the gorilla reference genome (**Table S6**). The western lowland gorilla sample A934_Delphi is not included in this study since it contains low-level contamination from a bonobo (Prado-Martinez et al. 2013). Consistent with this potential contamination, A934_Delphi shows an extreme value for the D-statistic relative to other western gorillas; however, significant statistics are also obtained when using other samples (**Table S6A**). We do not observe significant D-statistics for genotypes calculated from reads mapped to the gorilla reference genome using BSNP (**Table S6B**). Additional Cross River samples, as well as new analytic approaches that take advantage of the additional information contained in physically phased genome sequences (Schiffels and Durbin 2014) may shed further light on patterns of gene flow among extant gorilla species.

### Western Gorilla Demographic Inference

Given the availability of 14 western lowland gorilla samples, we estimated a single-population demographic history using *∂α∂i*. Due to limited sample size, our model does not incorporate other subspecies/species. Our *∂α∂i* analysis indicates that western lowland gorillas have undergone a small, ancient population size expansion event 970 kya followed by a drastic size reduction 23 kya. These results are broadly concordant with previous estimates of temporal population size change in gorillas based on the PSMC model (Prado-Martinez et al. 2013) (**Figure S14**), especially given that it is known that PSMC tends to smooth instantaneous size changes. We note that the ancient increase predates our estimation for the separation of eastern and western gorillas, and the recent size decrease post-dates our estimation of Cross River – western lowland separation. The underlying causes of these effective population size changes are unclear. Previous studies note glacial and interglacial oscillations during the last two million years may have had an effect on gorilla population size and structure (Thalmann et al. 2007). For example, during the Last Glacial Maximum, rainforest cover was greatly diminished, especially in West Africa where a few refugia were surrounded by tropical grassland (Jolly et al. 1997).

Previous studies suggest substructure within the western lowland gorilla species (Clifford et al. 2004; Fünfstück et al. 2014; Nsubuga et al. 2010, Scally et al. 2013), but our results support the use of a one-population model of western lowland gorillas (though there may be some subtle isolation-by-distance or demic structure). Earlier studies that involved analysis of SSR motifs (DNA microsatellites) provided some indications of substructure within western lowland gorillas (Fünfstück et al. 2014; Nsubuga et al. 2010). These results are not necessarily at odds with our analyses as the mutation rates vary significantly between genome-wide SNPs and microsatellites. While a slower evolving set of markers, such as SNPs, can identify expansion from a common ancestor and imply demographic changes over tens of thousands of generations, more rapidly evolving microsatellite loci can reveal more recent aspects of gene flow and population substructure. The gorillas utilized in this study have diverse origins; however, some origins cannot be precisely confirmed. PCA and ADMIXTURE analysis support grouping of samples into one population for *∂α∂i* analysis. Additionally, models inferred separately on subsets of the data yielded concordant results (**Table S2**).

In addition to the inferred decline in gorilla effective population size, census estimates note that the gorilla population has declined by more than 60% in the past 20-25 years, prompting their “critically endangered” conservation status (IUCN 2013). This decrease is thought to be due predominantly to Ebola outbreaks and commercial hunting (Walsh et al. 2003, Le Gouar et al. 2009). This sharp decline is much too recent to be observed in our analysis given the dataset available.

### Natural Selection in Western Lowland Gorillas Identifying Recent Selective Sweeps

Few previous studies have analyzed broad patterns of natural selection in the gorilla genome. Scally et al. (2012) found that regions exhibiting accelerated evolution in gorillas, compared to humans and chimpanzees, were most enriched for developmental terms including: ear, hair follicle, gonad and brain development, and sensory perception of sound. Our analysis complements these results by inferring regions with significant recent signs of selective sweeps within population-level full-genome western lowland gorilla data. Gene ontology enrichment analysis identified sensory perception of taste (GO:0050909) as one of the most significantly enriched category in the genome. This term has also been identified as enriched in selected genomic regions in mammalian genomes (Kosiol et al. 2008). All identified genes (*GOGO-T2R14*, *TAS2R19, TAS2R20*, *TAS2R50*) are type 2 taste receptors, which are thought to be responsible for bitter taste perception in humans (Adler et al. 2000; Chandrashekar et al. 2000; Matsunami et al. 2000). Bitter taste receptors are thought to be important to avoiding harmful substances, and have been predicted to have undergone an extensive gene expansion in mammalian evolution (Go 2006). Furthermore, among the top 16 most enriched GO terms are terms involving cardiac muscle function and fibroblast apoptosis. Interestingly, cardiomyopathy involving fibrotic proliferation is a prominent cause of death in captive gorillas, particularly in males (Schulman et al. 1995).

### Distribution of Fitness Effects

We found the distribution of fitness effects and the rate of fixation of adaptive mutations to be similar, but lower than estimates in humans (Boyko et al. 2008). This result is counterintuitive given that western lowland gorillas have a larger effective population size than humans. Because the mean gamma (N_e_*s*) is lower in gorillas and N_e_ is higher, we infer the magnitude of E(s) to be quite a bit smaller in gorillas than humans. Hvilsom et al. (2012) estimated the proportion of adaptive mutations driven to fixation in chimpanzee autosomes to not be significantly different from zero (95% CI: −0.09 – 0.07), which strongly overlaps with our estimates in western lowland gorillas. Our best-fit demographic model is a three-epoch model. To estimate the proportion of adaptive non-synonymous substitutions in the genome, DFE-alpha utilizes a two epoch demographic model. Messer and Petrov (2013) have previously shown that while the approach invoked by DFE-alpha generally correctly recovers *α*, Veeramah and colleagues (2014) demonstrated that DFE-alpha can substantially underestimate the true N_e_*s* because of background selection acting at linked sites. In addition the strength of any selection, gamma, acting at synonymous sites, which are taken as putatively neutral in this approach, is likely to be larger in gorillas due to their larger effective population size, potentially further distorting estimates of the DFE. As such while our estimate of *α* may be quite robust, the reliability of the DFE estimate is more uncertain.

### Conservation Implications

Conservation of wild gorilla populations in their habitats will benefit from focused efforts to protect populations that collectively encompass the genetic diversity of each species and subspecies. Identification of gene flow that occurred in the past between populations provides impetus for landscape-level conservation plans to provide for migration corridors inferred from genetic data. The evidence for gene flow between western lowland gorillas and eastern lowland gorillas, which have also been shown to have relatively low levels of genetic diversity, suggests a role of rare migration events in population sustainability, as seen in other species (Grant and Grant 2011). Managed populations of gorillas in zoos benefit from veterinary care that, increasingly, may benefit from medical approaches based on genetic information. Cardiac disease is a major mortality factor in managed gorilla populations (McManamon and Lowenstine 2011). The opportunities to provide supportive care based on an understanding of the evolutionary similarities and differences in cardiac development and physiology between gorillas and humans can contribute to the welfare of managed gorilla populations, while also providing insights into the evolution of loci associated with cardiac disease risk in humans.

## Material and Methods

### Samples

Samples without evidence of sequence read contamination from unrelated western lowland gorillas (n=14), eastern lowland gorillas (n=2) and a Cross River gorilla (n=1) were mostly obtained from blood from wild-caught zoo specimens (Table S1) (Prado-Martinez et al. 2013). All samples were sequenced on an Illumina sequencing platform (HiSeq 2000) with data production at three different sequencing centers; samples were sequenced to 12.7 - 42.1x coverage. Samples were collected under the supervision of ethical committees and CITES permissions were obtained as necessary. Sequence reads are available from the SRA under accession SRP018689.

### Mapping to Gorilla Reference Assembly

Sequences were mapped to gorGor3 and filtered as detailed in Prado-Martinez et al. (2013). Variants were identified in three pools of samples: the 14 western lowland gorillas, the 2 eastern gorillas, and the 1 Cross River gorilla sample. To compare variant calls among sample sets, we generated genome masks that identified all sites that were callable across all samples. Filters were calibrated such that we captured 90% of sites that passed the VQSR procedure (Prado-Martinez et al. 2013). For western lowland gorillas, the filters correspond to a total sample read depth (DP) >= 95 and <=307, mapping quality (MQ) >= 39 and percent of reads with mapping quality 0 (MQ0fraction) <= 3. For eastern lowland gorilla, the criteria were DP >= 12 and <=37, MQ >= 33, and MQ0fraction <=4. For the Cross River gorilla, the criteria were DP >= 5 and <=24, MQ >= 38 and MQ0fraction <=0. For each sample set, we additionally removed sites within 5 bp of called indels, and removed all positions overlapping with segmental duplications (Sudmant et al. 2013). For analysis of the site frequency spectrum, we additionally imposed a minimum depth criteria of eight to increase accuracy at singleton sites. For G-PhoCS analysis, variants were identified for each sample independently using BSNP to avoid bias induced by the reference genome and from population level genotype calling. The genotype coordinates were then converted from gorGor3 (Ensembl release 62) to gorGor3.1 (Ensembl release 64) using a custom script.

### Recombination rate estimates

Gorilla-specific recombination rates were estimated using western gorilla SNP data; described in detail in Stevison et al. (in prep). Briefly, using both the human-based mapping and the species-specific mapping described above, the data were filtered using a combination of vcftools (Danecek et al. 2011) and custom scripts. Sites with more than 80% missing data were removed. Then, variable sites within 15bp of each other were thinned to only retain a single site. Next, a reciprocal liftOver (minMatch=0.1) (Hinrichs et al. 2006) was performed to remove sites that did not map back to the original position. Finally, sites not in Hardy-Weinberg were removed (cutoff=0.001). After these initial filters were performed on the sites mapped to both reference genomes, the remaining sites were intersected between the two assemblies, with only the species-specific orientation used for subsequent phasing and rate estimation steps. Next, synteny blocks were defined based on the coordinates in both the human and non-human primate reference genomes. Then, within each syntenic region, phasing and imputation was performed using the software fastPHASE (Scheet and Stephens 2006), and an additional filter based on minor allele frequency was performed (cutoff=0.05). For improved phasing accuracy, the variants were re-phased using the software PHASE (Stephens and Donnelly 2003) similar to Auton et al. 2012. Rates were then estimated in 4000 SNP blocks using LDhat (Fearnhead and Donnelly 2001, The International HapMap Consortium 2005) (same run parameters as in Auton et al. 2012). The final number of sites used to estimate recombination rates was ∼7.8 million, as compared to 5.3 and 1.6 million for western chimpanzee and HapMap, respectively.

### Summary Measures: PCA, Population structure, Heterozygosity

Inference of population structure and principle components analysis, which require a set of independent SNPs, were conducted on 10% thinned data when comparing all three subspecies, using ADMIXTURE (Alexander et al. 2009) and smartpca (Patterson et al. 2006), respectively. PCA of three species was conducted on the intersect of the 8x data in western lowland, Cross River, and eastern lowland gorillas. When considering only the western lowland gorillas, data was pruned for linkage (plink –indep 50 5 2). We performed 10 independent ADMIXTURE runs for each tested value of K. Heterozygosity was estimated based on the number of heterozygous SNPs per individual in the unfiltered 8x data.

### Evolutionary Relationship of Western Lowland, Cross River and Eastern Lowland Gorillas

G-PhoCS utilizes input alignments from multiple independent “neutral loci” in which recombination within loci occurred at negligible rate but recombination between loci were sufficient to assume that genealogies are approximately uncorrelated (Gronau et al. 2011). Assuming that parameters of recombination are broadly consisted among primates, we adopted the 37,574 neutral loci previously identified by Gronau et al. (2011) for the human genome (build NCBI36), lifted-over these loci to the gorilla genome and then applied a series of filters to obtain a new set of “neutral loci” for the gorilla genome. Specifically, we removed regions without conserved synteny in human-gorilla alignments, recent transposable elements annotated by RepeatMasker with <=20% divergence, exons of protein-coding genes, conserved noncoding elements according to phastCons, and recent segmental duplications in Gorilla. This resulted in 26,248 loci, with size of ∼1 kb and interlocus distance of ∼50 kb. We called genotypes from the whole genome data at these neutral loci using BSNP, setting –P flat, which assumes uniform prior distribution to determine genotype calls for each individual without bias introduced by the reference genome. For each locus, we also masked simple repeats, positions within 3 bp of an insertion/deletion, positions with less than 5 reads, and CpG sites. Finally we used MUSCLE (Edgar 2004) to make alignments of each inferred sequence. After removing loci with completely missing data (all Ns) in at least one individual, we obtained a final set of 25,573 neutral loci for input to G-PhoCS.

We applied G-PhoCS to different combinations of samples. These combinations always included both eastern lowland gorillas, the single Cross River gorilla but contained different combinations of two western lowland gorillas. An aligned human reference genome was included as an outgroup. We first evaluated four alternative scenarios: no migration between gorillas species and bi-directional migration between any two gorillas species. For each case, we ran G-PhoCS for 50,000 iterations and found that this was sufficient to establish convergence for the no migration and bi-directional migration models between western and Cross River gorilla. We reran the analysis with bi-directional migration between western and eastern gorilla, and allowed two migration band parameters, one from western lowland to eastern lowland gorilla and another from Cross River to eastern lowland gorilla. We found that 300,000 iterations were sufficient to establish convergence for parameters of interests and we set the burn-in as the first two-thirds of iterations (**Figure S9**). The raw estimates by G-PhoCS are ratios between model parameters. Using humans as an outgroup, we calibrated the model based on the average genomic divergence time between human and gorilla, denoted T_div_. We assume a range of T_div_=8.0-12.0 Mya. An average mutation rate was calculated by 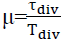. This mutation rate differs from the rate used in *∂α∂i* analyses because it ignores CpG mutations, which are excluded by our filters. Effective population sizes were calibrated by a factor of (4*19*µμ)^−1^, assuming an average gorilla generation time of 19 years (Langergraber et al. 2012). We also calculated estimates of expected number of migrants per generation, given by m_AB_ * *θ*_*B*_ and the total migration rate, given by m_AB_ * τ_AB_.

### Demographic Inference of Western Lowland Gorilla

The western lowland gorilla single population demographic model was inferred through a diffusion approximation approach implemented in the *∂α∂i* software (Gutenkunst et al. 2009). This approach calculates the log likelihood of the model fit based on a comparison between the observed and expected site frequency spectrum (SFS). Five demographic models were evaluated: a standard neutral model, an exponential growth model, a model of a bottleneck followed by exponential growth, a two epoch model and a three epoch model. We evaluated results with all SNPs that passed the filters and had at least 8x coverage, as well as a subset of these SNPs thinned to 100kb. For each model, ten independent runs were performed and the model and associated parameters that maximized the likelihood were chosen. To convert *∂α∂i* parameter output to years and effective population sizes, we assumed a mutation rate of 1.1 × 10^−8^ per generation (Roach et al. 2010; The 1000 Genomes Project Consortium 2010) and a generation time of 19 years (Langergraber et al. 2012). Confidence intervals for each parameter were determined through bootstrapping the input SNPs in blocks of 500 kb 1000 times.

For analysis of western lowland gorillas we used all sites from the genomic data with at least 8x coverage in all samples. The unfolded (polarized) SFS was determined using humans as an outgroup and ancestral misidentification was corrected using the method developed in Hernandez et al. (2007), which is implemented in *∂α∂i*. Briefly, this approach infers the unfolded SFS through a context dependent mutation model. It considers the trinucleotide sequence context of each SNP in gorillas and the outgroup, the great ape transition rate matrix for each nucleotide (as in Hwang and Green 2004, provided by Hwang DG, unpublished), the proportion of each trinucleotide sequence in the gorilla sequence data, and the gorilla-outgroup divergence (empirically estimated at 1.60% and 1.51% from the complete and 8x filtered sequence data, respectively).

### Signals of recent selective sweeps in Western Lowland Gorillas

Signals of recent selective sweeps were inferred using the SweepFinder method developed in Nielsen et al. (2005) implemented in SweeD (Pavlidis et al. 2013). The method uses a non-overlapping sliding window approach to calculate the composite likelihood of the data for two models: (1) a model of a recently completed selective sweep in the window and (2) a model that the window SFS is from the same distribution as the background SFS, where the background SFS is the SFS of the entire chromosome. This method outputs a composite likelihood ratio (CLR) of these two models. Non-overlapping windows of 100 kb along each chromosome were used to analyze the polarized 8x data. Windows that had less than 10% of the base pairs callable in the 8x dataset were excluded from analysis.

The unfolded SFS was determined using a two outgroup approach. Nucleotides at each SNP in the gorilla genome were compared with the reference alleles for human (hg19) and rhesus macaque (rheMac2). At each SNP position, if one gorilla allele matched the reference allele in both humans and rhesus macaques, then that allele was assumed to be the ancestral allele. If the three species did not share an allele at a specific SNP, the site was excluded. The ancestral misidentification correction implemented in Hernandez et al. (2007) adjusts the overall SFS and not individual SNPs, and was therefore inappropriate for this analysis.

To determine CLR significance, neutral genomic regions were simulated with ms (Hudson 2002). This test may be weakly dependent on demography, recombination rate, and “callable” sequence length (Nielsen et al. 2005, Williamson et al. 2008); therefore, all neutral simulations are based on the inferred three epoch demography, as well as conservative estimates of the recombination rate and “callable” sequence length. Though the CLR is rather robust to variable recombination rates, regions of low recombination may be more likely to look like selective sweeps (Nielsen et al. 2005). Furthermore, regions with less “callable” sequence have less data and thus may have lower power to recognize a selective sweep through the CLR. Due to this, 100,000 neutral regions were simulated for each of the following recombination rates (in cM/MB): 0, 0.25, 0.5, 1, and 2. In each region, base pairs were randomly masked at one of the following levels: 90%, 80%, 70%, 60%, 50%. A total of 2,500,000 regions were simulated; 100,000 for each combination of parameters. The average recombination rate in each gorilla 100 kb genomic region was calculated from a gorilla specific recombination map (Stevison et al., in prep). For gorilla regions without an estimated recombination rate, the average recombination rate (0.6429 cM/Mb) was used (Stevison et al., in prep). The CLR significance of each gorilla 100 kb region was determined through comparison with the closest set of neutral simulations. When gorilla windows were between parameters, simulations with lower recombination and higher masking were used, making the test more conservative.

We utilized FDR methods outlined in Storey and Tibsirani (2003) and the tuning parameter selection method from Williamson et al. (2007) to estimate the percent of features we call “significant” that are actually null. The tuning parameter selection method was used because, as Williamson et al. (2007) points out, the CLR was designed to be conservative and thus there are many regions with p=1. This violates Storey and Tibsirani’s (2003) assumption that p-values of null features follow a uniform distribution. As we are testing many hypotheses simultaneously, we estimated the proportion of inferred selected genomic regions likely to be false positives at various p-values thresholds. We utilized the approach by outlined in Storey and Tishirani (2003) and the tuning parameter selection method from Williamson et al. (2007). Using the same parameters as Williamson et al. (2007), we found that the FDR at a p-value threshold of 10^−5^ was 0.50%, at 10^−4^ was 3.4%, at 0.001 was 9.11%, and at 0.01 was 34.04%.

Genomic features, including genes and their corresponding GO terms, in regions with significant signs of selective sweeps were identified through the Ensembl database (Flicek et al. 2011). All genes reported are verified human genes that are computationally predicted to have an orthologous gene in the gorilla genome. The Bioconductor package, topGO, was used for gene ontology enrichment analysis (Alexa and Rahnenfuhrer 2010). The set of significant genes tested has p < 0.001 and the background distribution of genes were those that overlapped with windows tested in SweeD (regions with > 10% of their sequence in the callable genome). The elimination method with Fisher’s exact test was used to infer significantly enriched GO terms.

### Distribution of fitness effects

The distribution of fitness effects was inferred through the DFE (distribution of fitness effects)-alpha server (Keightley and Eyre-Walker 2012). This method attempts to correct for the biases in the McDonald-Kreitman test due to slightly deleterious mutations. Briefly, this method simultaneously infers demography and the distribution of fitness effects, based on transition matrix methods. Adaptive substitutions are inferred through the difference between the observed divergence and the predicted divergence, based on a gamma distribution and a two epoch demographic model. Input data was the folded nonsynonymous and synonymous frequency spectra, as annotated by SNPEff (Cingolani et al. 2012). To calculate the number of divergent sites, we utilized the UCSC multiz alignment of the human (hg19), chimpanzee (panTro3), and rhesus macaque (rheMac2) genomes to the gorilla (gorGor3) genome coding region and restricted to sites with no missing data. The number of divergence sites was then calculated from the sites where the human, chimpanzee, and rhesus macaque shared the same allele, and the gorilla genome allele differed. Confidence intervals were determined through bootstrapping the input synonymous and nonsynonymous SNPs 1000 times.

## Supplementary Material

Supplementary tables S1-S5 and figures S1-S13.

## Acknowledgements

The authors thank Ryan Gutenkunst for assistance with *∂α∂i* and Omar Cornejo for extensive input on analytical methods. This work was supported by the National Institutes of Health to MFH. and JDW. (R01_HG005226). KFM is supported by the National Institutes of Health (2T32GM007276-­-39) and a Stanford Center for Computational, Evolutionary and Human Genomics (CEHG) fellowship. AEW is supported by National Science Foundation Graduate Research Fellowship Grant DGE-1143953.

